# CRISPR-Cas9 knockout screen informs efficient reduction of the *Komagataella phaffii* secretome

**DOI:** 10.1101/2024.04.21.590449

**Authors:** Neil C. Dalvie, Timothy R. Lorgeree, Yuchen Yang, Sergio A. Rodriguez-Aponte, Charles A. Whittaker, Joshua A. Hinckley, John J. Clark, Amanda M. Del Rosario, Kerry R. Love, J. Christopher Love

## Abstract

The yeast *Komagataella phaffii* is widely used for manufacturing recombinant proteins, but secreted titers of recombinant proteins could be improved by genetic engineering. In this study, we hypothesized that cellular resources could be redirected from production of endogenous proteins to production of recombinant proteins be deletion of unneeded endogenous proteins. We identified a set of endogenous secreted proteins in *K. phaffii* and attempted to disrupt these genes, but our efforts were hindered by limited annotation of genes, especially essential ones—this is a common problem for genetic engineering of non-model organisms. To predict essential genes, therefore, we designed, transformed, and sequenced a pooled library of guide RNAs for CRISPR-Cas9-mediated knockout of all endogenous secreted proteins. We then used predicted gene essentiality to guide iterative disruptions of up to 11 non-essential genes. Engineered strains exhibited a ∼20x increase in the production of human serum albumin and a twofold increase in the production of a monoclonal antibody. The pooled library of secretome-targeted guides for CRISPR-Cas9 and knowledge of gene essentiality reported here will facilitate future efforts to engineer *K. phaffii* for production of other recombinant proteins and enzymes.

## Introduction

There is growing interest in alternative microbial hosts as manufacturing chassis to produce recombinant proteins,^1,2^ including ones with therapeutic uses typically manufactured in mammalian cells. The methylotrophic yeast *Komagataella phaffii* (*Pichia pastoris*) offers unique advantages compared to the conventional model microorganisms *Escherichia coli* and *Saccharomyces cerevisiae* because of its productive secretory pathway.^3–5^ *K. phaffii* is routinely used for large-scale manufacture of small therapeutic proteins (<30 kD) such as insulin,^6^ vaccine antigens,^7^ and VHH antibodies.^8^ In addition, *K. phaffii* has now been used for the commercial production of a full length monoclonal antibody (mAb) as well (eptinezumab).^9^

Mammalian cell lines such as Chinese hamster ovary (CHO) and human embryonic kidney (HEK293) have required several decades of empirical selections and process-related optimizations to manufacture mAbs and other large proteins efficiently and reliably.^10^ Emerging applications of gene editing in CHO cells have shown that the knockout of up to 14 natively secreted host cell proteins (HCPs) can improve both the secreted titer and purity of a recombinant mAb.^11^ In contrast to CHO cells, *K. phaffii* secretes a limited number of HCPs,^12^ which can result in high initial purity of recombinant proteins in culture supernatant and facilitate purification and characterization of the product.^13^ Proteins secreted at lower titers, however, may compete with HCPs for cellular resources including amino acids, ribosomes, protein folding machinery, and secretory capacity.^14–16^ We hypothesized that knockout of natively secreted proteins may improve recombinant protein secretion in *K. phaffii*.

Here, we characterized a set of proteins from *K. phaffii* identified in the culture fluids after fermentation,and disrupted up to 11 genes that code for these secreted proteins. Several engineered strains, especially one with six disrupted genes, exhibited improved production of multiple large (>50 kDa) human proteins. To facilitate the iterative knockout of more than three secreted proteins, we performed a pooled CRISPR-Cas9 knockout library to measure the essentiality of all secreted proteins. This knowledge of gene essentiality and new capability for pooled screening methods should also inform future efforts to engineer other cellular processes or pathways in *K. phaffii* for improved production of recombinant proteins.

## Results

### Identification and knockout of secreted proteins

We sought to identify the set of proteins manifest in the culture fluids during fermentation (the secretome) in *K. phaffii*. We computationally identified 257 coding sequences in the *K. phaffii* genome with putative secretory signal peptides (Methods, Table S1). We then cultured wild type *K. phaffii* (NRRL Y-11430) and analyzed the proteins found in the extracellular fluid by mass spectrometry (Fig. S1).^17^ We detected 134 proteins (Table S1). The relative abundance of most proteins was similar between the glycerol and methanol medium (R = 0.93), respectively (Fig. 1A). (These two sources of carbon are commonly used to accumulate biomass and induce recombinant protein production, respectively.) Interestingly, only 30 of the proteins identified in the cell cultivation fluids by mass spectrometry had computationally predicted signal peptides. A significant number of these proteins were previously found to be enriched in microsomes (endoplasmic reticulum) (p<0.0001 in glucose medium, p<0.001 in methanol medium) and in the very early Golgi (p<0.01 in methanol medium) (Fig. S2), suggesting these proteins may be secreted by the canonical yeast protein secretory pathway.^18^ We did not detect 227 additional proteins predicted to contain signal peptides. We hypothesize that these proteins were present at concentrations too low to detect or are targeted to other cellular organelles (and therefore are not secreted). A significant number of these proteins were previously found to be enriched in organelles such as the very early Golgi, early Golgi, microsome, vacuole, and mitochondria, all of which use signal peptides for protein localization (Fig. S2).^19,20^ Finally, we experimentally detected 104 proteins not predicted to contain secretory signal peptides. A significant number of these proteins were associated with organelles such as the cytosol, mitochondria, and peroxisome (Fig. S2). These proteins were also previously found in most organelles and cell fractions, even if not statistically enriched (Fig. S3). The genes that code for these proteins are also highly expressed (p< 0.0001) (Fig. S4).^17^ We postulate that these proteins are abundant in the cell and may escape into the extracellular space by cell lysis or non-specific packaging into vesicles. These abundant intracellular proteins may also compete with the recombinant protein for cellular resources during transcription and translation.^14^ Based on this analysis, we defined the secretome of *K. phaffii*, therefore, as the collection of 361 proteins that were predicted to contain a signal peptide or that were detected in culture supernatants (Table S1).

**Fig. 1.**
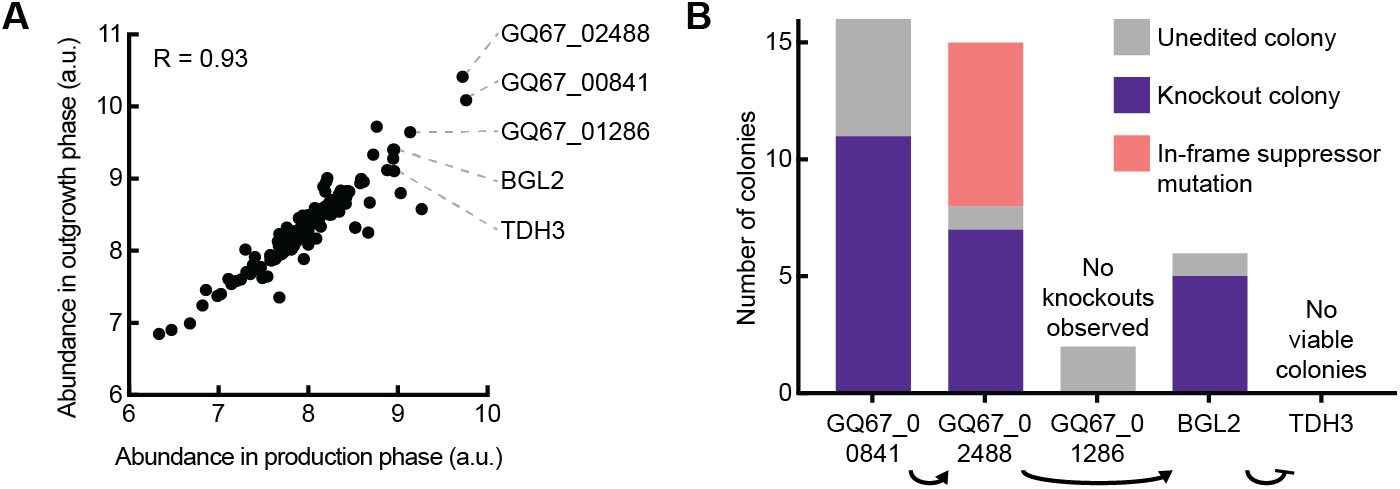
Identification and knockout of secreted proteins. A) Relative abundance of proteins detected in culture supernatant harvested from cultures in glycerol medium (outgrowth) or methanol medium (production). B) Knockout efficiencies of sequential disruption of the most abundant proteins in the *K. phaffii* secretome.

To improve the production of recombinant proteins, we next sought to disrupt the most abundant secreted proteins based on a rank-ordering of our initial analysis and using a previously reported, host-informed strategy for CRISPR-Cas9 genome editing in *K. phaffii*.^21,22^ We used this tool to serially disrupt several of the most abundant secreted proteins. We encountered engineering challenges in this approach, however, including in-frame deletions in disrupted genes, and were unable to disrupt two of the first five targeted genes (Fig. 1B). (GQ67_01286 is a homolog of the gene *ayr1* in *S. cerevisiae*—*ayr1* and *tdh3* are both non-essential in *S. cerevisiae* (Saccharomyces Genome Database).) We posited that identification of essential genes—particularly unannotated essential genes—would streamline further engineering of *K. phaffii*. We therefore sought to identify which genes among the identified secretome of *K. phaffii* are essential in laboratory conditions (under standard conditions for fermentation).

### Identification and knockout of non-essential secreted proteins

We aimed to evaluate the essentiality of the genes encoding the secretome of *K. phaffii* in parallel. One key innovation in our previously reported CRISPR-Cas9 tool was the reduction of the number of nucleotides that must be replaced in a single guide RNA (sgRNA) cassette to retarget cleavage of DNA by Cas9, enabling pooled synthesis of sgRNA libraries.^21^ We created a pooled library of sgRNAs for CRISPR-Cas9-mediated disruption of all genes in the secretome (Fig. 2A) and from the resulting screen, calculated an “essentiality” score for each one (see Supplementary Methods, Table S1). Genes that had the highest essentiality scores included ribosome subunits (*rpl16a*), translation factors (*tef1*), and essential enzymes (*tdh3/gapdh, pgk1*). We performed weighted gene set enrichment analysis (GSEA) on the essentiality scores of all 361 genes and identified several gene sets enriched with essential secretome genes including carbohydrate metabolism, translation, and membrane transport (Fig. 2B). These observations suggested that the screen with the secretome-directed library was successful for scoring the essentiality of those genes in *K. phaffii*.

**Fig. 2.**
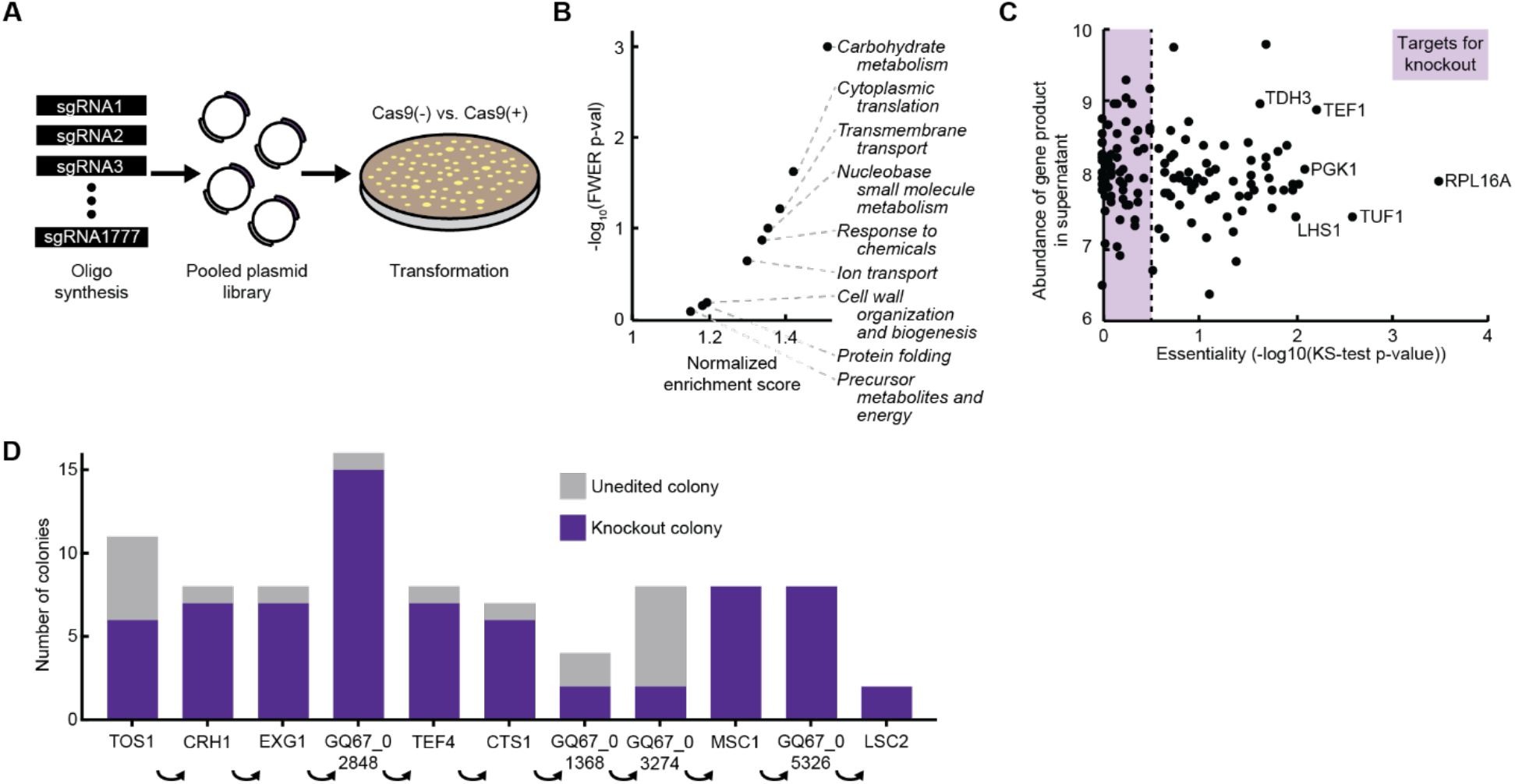
Identification and knockout of non-essential secreted proteins. A) Schematic of CRISPR-Cas9 knockout screen to determine essentiality. B) Gene set enrichment analysis of genes in the *K. phaffii* secretome weighted by their essentiality score. C) Plot of gene essentiality and relative abundance of proteins in production phase culture supernatant. Genes with an essentiality score of >0.5 were considered likely to be essential. D) Knockout efficiencies of sequential disruption of non-essential genes in the *K. phaffii* secretome. Knockout genotypes were determined by Sanger sequencing of the targeted locus for up to 16 colonies per transformation.

We next prioritized a short list of non-essential genes from the secretome as engineering targets. We filtered the secretome for 61 genes with an essentiality score less than 0.5 and with gene products experimentally detected in the culture supernatant (Fig. 2C). Then, we removed seven genes from the potential target pool known to contribute to methanol metabolism since disruption of these genes may affect cellular function during methanol-induced protein secretion (though appear non-essential when cultured on solid glucose medium). Finally, we constructed and separately transformed 14 CRISPR-Cas9 vectors with multiplexed sgRNAs targeting up to four genes per vector (Fig. S5). On the first attempt, we observed disruption of 20 of the 54 gene targets. We hypothesized that certain combinations of multiplexed knockouts may cause unforeseen synthetic lethality. It may be feasible to disrupt more of these genes with further optimization. We chose to proceed, however, with engineering strains based on the 20 gene disruptions observed.

We next attempted to combine many disruptions to create strains with a reduced secretome. We performed gene disruptions sequentially in two lineages (SΔ3a (*Δtos1, Δcrh1, Δexg1*) and SΔ3b (*Δtfs1, Δlsc2, Δgq67_05326*)), and then combined these sets to construct a new strain SΔ6 (*Δtos1, Δcrh1, Δexg1, Δtfs1, Δlsc2, Δgq67_05326*). We also extended the SΔ3a lineage to construct SΔ9 with six additional disruptions (*Δgq67_02848, Δtef4, Δcts1, Δgq67_01368, Δgq67_03274, Δmsc1*). We added Δ*lsc2* and Δ*gq67_05326* to SΔ9 to create SΔ11, but we were unable to disrupt *tfs1* in SΔ11. This observation suggested that *tfs1* may confer synthetic lethality with another disrupted gene in SΔ11.

Throughout this engineering process, we noticed high disruption efficiencies (typically 80-100% of colonies were disrupted). During construction of SΔ11, for example, we combined all 11 knockouts without screening more than 16 colonies at each step (Fig. 2D). We attributed this engineering efficiency to the additional knowledge of gene essentiality used to guide the selection of the targeted genes. We next sought to assess the utility of these engineered strains for production of recombinant proteins.

### Productivity and growth of secretome-deficient strains

We evaluated the secreted productivity of SΔ3a, SΔ3b, SΔ6, and SΔ11 compared to the parent strain (Table 1). We first transformed strains with a vector enabling the secreted expression of human serum albumin (HSA), a 67 kDa protein using the methanol-responsive promoter PAOX1. We cultivated cells in glycerol-containing medium to build biomass, induced expression of the recombinant gene by replacing the medium with a methanol-containing one for 24 hours, and evaluated the extracellular protein titer (Fig. 3A-B). Surprisingly, while neither strain with three knockouts exhibited a significant change in specific productivity, the strain with all six knockouts (SΔ6) exhibited a ∼20-fold increase in protein titer and specific productivity (protein titer normalized to the biomass of the culture based on measured optical density).

**Table 1.**
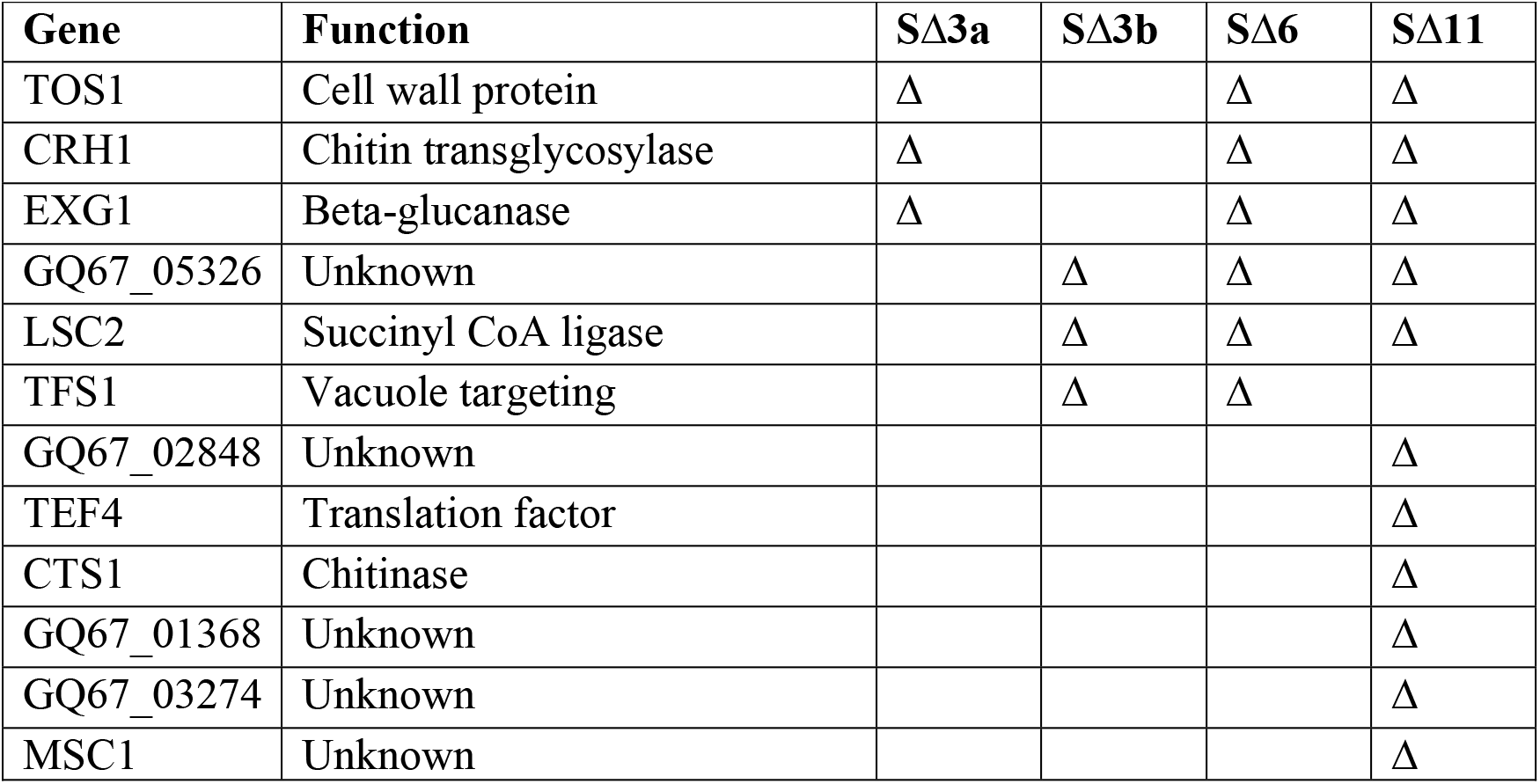
Engineered strains with reduced secreted proteins.

**Fig. 3.**
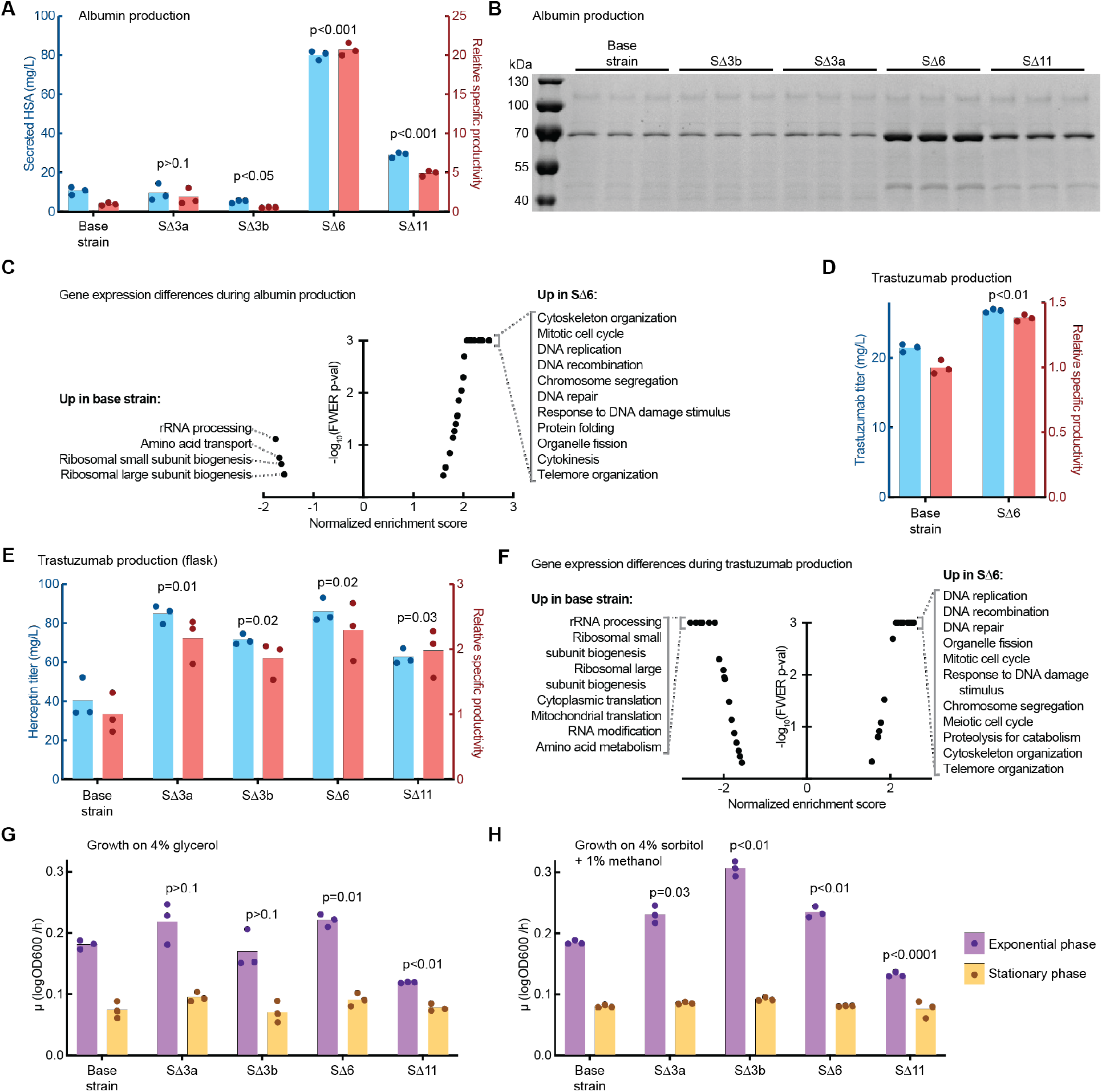
Productivity and growth of engineered knockout strains. A) Secreted titer and specific productivity of HSA from 3 mL microplate cultures. B) SDS-PAGE of 3 mL microplate culture supernatant. HSA protein is visible at ∼70 kDa. C) Enriched gene sets between SΔ6 and the base strain during production of albumin. D) Secreted titer and specific productivity of trastuzumab in 3 mL microplate cultures. E) Secreted titer and specific productivity of trastuzumab in 100 mL shake flask cultures. F) Enriched gene sets between SΔ6 and the base strain during production of trastuzumab. G) Growth rates of strains in 200 µL cultures in glycerol outgrowth medium. H) Growth rates of strains in 200 µL cultures in methanol production medium. In all bar plots, significance of specific productivity or exponential growth rate compared to the base strain was determined by unpaired Welch’s t-test.

To assess the differences between the SΔ6 and wildtype (control) strains producing HSA, we analyzed the transcriptional states of the cells by RNA-seq during recombinant protein expression. Genes related to ribosomal processing and translation were upregulated in the base strain, while genes related to cell division and genome replication were upregulated in SΔ6 (Fig. 3C, Table S2). We hypothesize that the engineering changes in SΔ6 may alleviate the translational burden experienced by the base strain, either by reduction of the overall translational load from knockout of abundant secreted proteins, or by specific functions of the disrupted genes.

Given the improved production of HSA, we hypothesized that SΔ6 may also have benefits for producing other large proteins (such as mAbs). We evaluated the secreted titer of trastuzumab, a mAb used to treat HER2+ breast cancer. SΔ6 exhibited a ∼30% increase in specific productivity compared to the wildtype strain in 3 mL cultures (p = 0.002, unpaired Welch’s t-test) (Fig. 3D). To evaluate the performance of engineered strains at higher cell densities, we cultivated the base strain and all four engineered strains producing trastuzumab in 100 mL cultures in shake flasks. At this scale, all strains reached an optical density of 40-50 OD600 after one day of production. In these growth conditions, all the engineered strains secreted two-fold more trastuzumab than the base strain, particularly SΔ6 and SΔ3a (Fig. 2E).

We also assessed the gene expression of SΔ6 and the base strain during expression of trastuzumab. Like the strains that produced HSA, we observed that genes related to ribosomal and RNA synthesis were upregulated in the base strain, while genes related to cell and genome replication were upregulated in SΔ6 (Fig. 2F, Table S2). An overall reduction of the translational load may also improve production of trastuzumab, similar to the results for producing HSA.

Finally, we measured the rate of growth by seeding engineered antibody-producing strains at low density in 200 µL cultures. Interestingly, in the glycerol-containing media used to accumulate biomass, SΔ6 exhibited a higher growth rate than the base strain while SΔ11 exhibited a lower growth rate (Fig. 2G). Similarly, in the methanol-containing media used to induce expression of the recombinant protein, we observed higher growth rates for SΔ3a, SΔ3b, and SΔ6, and a lower growth rate for SΔ11 (Fig. 2H). These results, together with the observed improvement in recombinant protein titers, demonstrate that strains of *K. phaffii* with a reduced secretome can improve the secreted productivity of multiple proteins relevant for biopharmaceutical and vaccine products without a decrease in growth rate compared to the base strain.

## Discussion

We engineered four new strains of *K. phaffii* with improved productivity of recombinant proteins. One strain in particular, SΔ6, exhibited large improvements in extracellular titer of HSA (∼20x) and trastuzumab (∼2x) without a reduction in growth rate. This strain showed reduced expression of genes related to translation and synthesis of ribosomes. SΔ6 may have an increased cellular capacity for translation of the recombinant protein due to less translational demand from the native proteome. We also observed increased growth rates by SΔ3a, SΔ3b, and SΔ6 during production of trastuzumab. Translational capacity or the availability of amino acids may represent a general limitation for yeasts during recombinant protein production, therefore.^23–25^ This hypothesis is corroborated by another engineered strain with an upregulated translation factor that exhibited with improved secreted productivity by expanding the cellular capacity for translation.^26^ Strategies to further redirect translational capacity towards the recombinant product of interest warrant further investigation.

The improved productivity observed with SΔ6 may also result from the functions of specific disrupted genes or combinations of disrupted genes.^27^ We did not perform comprehensive combinatorial studies to determine how each individual disrupted gene affects the secretion of recombinant proteins. The gene *tfs1*, disrupted in SΔ3b, SΔ6, and SΔ11, may improve secretion of recombinant proteins by reducing the amount of protein directed towards the vacuole—a common degradation pathway for heterologous proteins.^28,29^ Similarly, three genes disrupted in SΔ6 are involved in construction of the yeast cell wall. Cell wall proteins are abundantly secreted and may consume a large fraction of amino acid, translational, and secretory resources.^30^ Disruption of the physical cell wall may also facilitate diffusion of large proteins through the cell wall and into the extracellular space.^31,32^ Deeper understanding of the impact of vacuolar and cell wall-related genes on recombinant protein secretion may inform further engineering.

We also report here the first pooled CRISPR-Cas9 screen in *K. phaffii*. We used this screen to predict the essentially of all genes that code for secreted proteins. Knowledge of essential genes will enable strain engineers to avoid targeting unannotated essential genes, which disrupts workflows for genome editing. In *K. phaffii*, 108 of the 361 proteins in the secretome are described as hypothetical proteins (Table S1), and only 218 proteins in the secretome have homologs in the model yeast *S. cerevisiae*.^17,33^ With knowledge of gene essentiality, we successfully disrupted seven unannotated genes without additional effort or screening (Fig. S5) and successfully knocked out 11 genes in sequence before encountering synthetic lethality. The predicted gene essentiality documented here will facilitate engineering of other pathways and functions in *K. phaffii* without the need for further pooled screening.

Finally, the sgRNA library used here targeted only one gene per cell and thus was unable to predict synthetic interactions between disrupted genes that may reduce fitness. Indeed, the SΔ11 strain, with the most disrupted genes, exhibited reduced growth rates during production of trastuzumab. We previously demonstrated that the sgRNA library design used here is compatible with multiplexed gene editing, which will enable pairwise or higher multiplexed knockout libraries in the future.

High-throughput functional genomics tools such as transposon libraries, oligo-mediated recombineering, and Cas9-mediated knockout or upregulation libraries are widely applied to model hosts such as *E. coli* and *S. cerevisiae*, including multiplexed libraries.^34^ When paired with high-throughput screens or selections, pooled genetic libraries enable identification of genes and pathways that may be tractably engineered to impact the desired phenotype.^35^ Pooled screening would be especially useful in non-model microbial hosts in which the functions of many genes are unknown.^17,36,37^ The sgRNA library design described here leverages native host tRNAs, which makes this approach a general strategy for pooled screening in non-model microbial hosts for production of recombinant proteins such as *K. phaffii, Trichoderma reesei, Hansenula polymorpha*, and *Aspergillus oryzae*.^21,38^

## Materials and methods

### Yeast strain cultivation

Strains were grown in 3 mL cultures in 24-well deep well plates (25°C, 600 rpm) or 100 mL cultures in 500 mL shake flasks (25°C, 300 rpm). Cells were cultivated in complex media (potassium phosphate buffer pH 6.5, 1.34% nitrogen base w/o amino acids, 1% yeast extract, 2% peptone). Cells were inoculated at 0.1 OD600, outgrown for 24 h with 4% glycerol feed, pelleted, and resuspended in fresh media with 3% methanol for HSA production, or 1% methanol, 40 g/L sorbitol, and 10 mM glutathione for trastuzumab production. Supernatant samples were collected after 24 h of production and analyzed.

For quantitative measurement of secreted protein titer, strains were cultivated in biological triplicate from frozen stocks. For growth assays, strains that produce trastuzumab were seeded at an optical density of 0.01 OD600 in 200 µL cultures (25°C, 300 rpm) in either outgrowth or production media in biological triplicate from frozen stocks. OD600 was measured every hour for 48 h and plotted on a log axis. Growth rate was calculated manually as the slope of the curve during exponential or stationary growth.

### Yeast strain construction

All strains were derived from wild-type *Komagataella phaffii* (NRRL Y-11430). Strains for recombinant protein production were derived from a modified base strain (AltHost Research Consortium Strain S-63 (RCR2_D196E, RVB1_K8E)) described previously.^39^ Genes containing recombinant protein products HSA and trastuzumab were synthesized (Integrated DNA Technologies) and cloned into a custom vector with the methanol-inducible promoter PAOX1. To enable protein secretion, HSA was expressed with the signal peptide from the *S. cerevisiae* α-mating factor, and trastuzumab was expressed with the signal peptide from the *S. cerevisiae* α-mating factor for the light chain and the signal peptide from human HSA for the heavy chain. All vector sequences are listed in the Supplemental Materials.

*K. phaffii* strains were transformed as described previously.^21^ After transformation of the HSA or trastuzumab expression vectors, 4-8 clones were selected and grown in 3 mL cultures. Supernatant samples were analyzed by SDS-PAGE, and the clone that exhibited the highest productivity was selected for quantitative growth and titer measurements.

Knockout of individual genes was performed with a custom knockout cassette as described previously.^22^ Disruption of genes was confirmed by PCR and Sanger sequencing. Design, construction, and screening of the pooled knockout library is described in the Supplemental Methods.

### Analytical assays for protein characterization

SDS-PAGE was carried out as described previously.^40^ HSA supernatant titers were measured by reverse phase liquid chromatography. Trastuzumab supernatant titers were measured by Protein A biolayer interferometry. Specific productivity was calculated as titer normalized to cell density by OD600, relative to the original base strain.

### LCMS measurement of the K. phaffii secretome

Wild-type *K. phaffii* was cultivated in 200 mL shake flask cultures in complex media. Cells were inoculated at 0.1 OD600, outgrown for 48 h with 4% glycerol feed, pelleted, and resuspended in fresh media with 1.5% methanol feed to simulate recombinant gene expression. Supernatant samples were collected after each phase of the cultivation.

Supernatant was reduced (10 mM dithiothreitol, 56 °C for 45 min) and alkylated (50 mM iodoacetamide, room temperature in the dark for 1 h). Proteins were subsequently digested with trypsin (sequencing grade, Promega, Madison, WI), at an enzyme/substrate ratio of 1:50, at room temperature overnight in 100 mM ammonium acetate pH 8.9. Trypsin activity was quenched by adding formic acid to a final concentration of 5%. Peptides were desalted using C18 SpinTips (Protea, Morgantown, WV), lyophilized, and stored at −80 °C.

Peptides were labeled with TMT 6plex (Thermo) per manufacturer’s instructions. Lyophilized samples were dissolved in 70 μL ethanol and 30 μl of 500 mM triethylammonium bicarbonate (pH 8.5), and the TMT reagent was dissolved in 30 μl of anhydrous acetonitrile. The solution containing peptides and TMT reagent was vortexed and incubated at room temperature for 1 h. Samples labeled with the ten different isotopic TMT reagents were combined and concentrated to completion in a vacuum centrifuge. The samples were labeled using the TMT 10plex channels as follows: 126 – 4/27/16 48 h induction; 127N – 4/29/16 48 h harvest; 127C – 5/6/16 96 h harvest; 128N – 5/4/16 48 h induction; 129N – 5/13/16 96 h harvest; 129C – 5/20/16 96 h harvest; 130N – 5/18/16 48 h induction; 130C – 5/27/16 96 h harvest; 131 –5/25/16 48 h induction.

Peptides were loaded on a precolumn and separated by reverse phase HPLC (Thermo Easy nLC1000) over a 140-minute gradient before nanoelectrospray using a QExactive Plus mass spectrometer (Thermo). The mass spectrometer was operated in a data-dependent mode. The parameters for the full scan MS were: resolution of 70,000 across 350-2000 *m/z*, AGC 3e^6^, and maximum IT 50 ms. The full MS scan was followed by MS/MS for the top 10 precursor ions in each cycle with a NCE of 34 and dynamic exclusion of 30 s. Raw mass spectral data files were searched using Proteome Discoverer (Thermo) and Mascot version 2.4.1 (Matrix Science). Mascot search parameters were: 10 ppm mass tolerance for precursor ions; 15 mmu for fragment ion mass tolerance; 2 missed cleavages of trypsin; fixed modification were carbamidomethylation of cysteine and TMT 10-plex modification of lysines and peptide N-termini; variable modification was methionine oxidation. TMT quantification was obtained using Proteome Discoverer and isotopically corrected per manufacturer’s instructions. Only peptides with a Mascot score greater than or equal to 25 and an isolation interference less than or equal to 30 were included in the quantitative data analysis. Relative abundance of each protein was defined by the log10 of the total area under the curve for all peptide counts detected for each protein, summed over four independent replicate cultivations.

### Analysis of the K. phaffii secretome

We determined which genes in the *K. phaffii* genome contained a signal peptide using SignalP 5.0 and filtering for Sec/SPI > 0.5.^41^ Comparison of the *K. phaffii* secretome to Valli et al.^18^ was performed by manual comparison in SnapGene (snapgene.com) of protein coding sequences from both the Love et al. genome^17^ and the genome from Pichiagenome.org.^42^ Descriptions of protein functions were obtained using BLAST.

### Transcriptome analysis

Cells were cultivated at 3 mL plate scale and harvested after 18 h of production in methanol medium. RNA was extracted and purified according to the Qiagen RNeasy 96 kit. RNA quality was analyzed on an Agilent BioAnalyzer to ensure RNA Quality Number >6.5. RNA was reverse transcribed with Superscript III (ThermoFisher) and amplified with KAPA HiFi HotStart ReadyMix (Roche). RNA libraries were prepared using the Nextera XT DNA Library Preparation Kit with the Illumina DNA/RNA UD Indexes Set A. sequenced on an Illumina Nextseq to generate paired reads of 50 (read 1) and 50 bp (read 2). Sequenced mRNA transcripts were demultiplexed using sample barcodes, aligned to the WT.fa Komagataella phaffii genome (strain Y11430) and exogenous transgenes, and quantified using Salmon version 1.6.0.^43^ Gene level summaries were prepared using tximport version 1.24.0^44^ running under R version 4.2.1.^45^ Gene set enrichment analysis (GSEA) was performed with GSEA 4.1.0 using Wald statistics calculated by DESeq2^46^ and gene sets from yeast GO Slim.^47^

## Acknowledgements

This work was supported by the AltHost Research Consortium. The content is solely the responsibility of the authors and does not necessarily represent the official views of the members of the AltHost Consortium. This work was partially supported by Cancer Center Support (core) Grant P30-CA14051 from the NCI to the Barbara K. Ostrom (1978) Bioinformatics and Computing Core Facility of the Swanson Biotechnology Center. N.C.D. was supported by a graduate fellowship from the Ludwig Center at MIT’s Koch Institute. J.A.H. was supported by a postdoctoral fellowship from the Ludwig Center at MIT’s Koch Institute.

## Author contributions

N.C.D., K.R.L., and J.C.L. developed the concepts and designed the study. J.J.C., A.D.R, and K.R.L. performed secretome characterization. N.C.D. and C.A.W. designed CRISPR libraries. N.C.D., T.L., and Y.Y. engineered yeast strains. S.R.A. and N.C.D. performed strain and protein characterization. N.C.D., C.A.W., and J.A.H. performed transcriptomics. N.C.D. and J.C.L. wrote the manuscript.

## Data availability

Plasmid sequences are included in the Supplemental Materials. Raw data from design and analysis of the pooled DNA library is included in the Supplemental Materials. Raw transcriptomic data is available on the NCBI Gene Expression Omnibus (accession number: GSE252605).

