## Supplement for "CRISPR-Cas9 knockout screen informs efficient reduction of the *Komagataella phaffii* secretome": Supplemental Materials.pdf

**Supplemental Material**

Neil C. Dalvie[1,2], Timothy M. Lorgeree[2], Yuchen Yang[1,2], Sergio A. Rodriguez-Aponte[2,3], Charles A. Whittaker[2], Joshua A. Hinckley[2], John J. Clark[2], Amanda del Rosario[2], Kerry R. Love[1,2], J. Christopher Love[1,2]

[1]Department of Chemical Engineering, Massachusetts Institute of Technology, Cambridge, Massachusetts 02139, United States
[2]The Koch Institute for Integrative Cancer Research, Massachusetts Institute of Technology, Cambridge, Massachusetts 01239, United States
[3]Department of Biological Engineering, Massachusetts Institute of Technology, Cambridge, Massachusetts 02139, United States

**Supplemental files:**

Supplemental Methods (for CRISPR library generation and screening)
R-code for sgRNA KS-test

**Tables:**

Table S1 – Lists and data for secretome gene selection and essentiality
Table S2 – Raw data from Gene Set Enrichment Analysis of engineered strains
Table S3 – Lists and data for sgRNA library generation, amplicon sequencing, and analysis

**DNA sequences attached:**

Vector for sgRNA library cloning (pND389)
Vector for Cas9 integration into the genome (pND386)
Vector for expression of HSA (D-17)
Vector for expression of trastuzumab light chain (D-67)
Vector for expression of trastuzumab heavy chain (D-85)

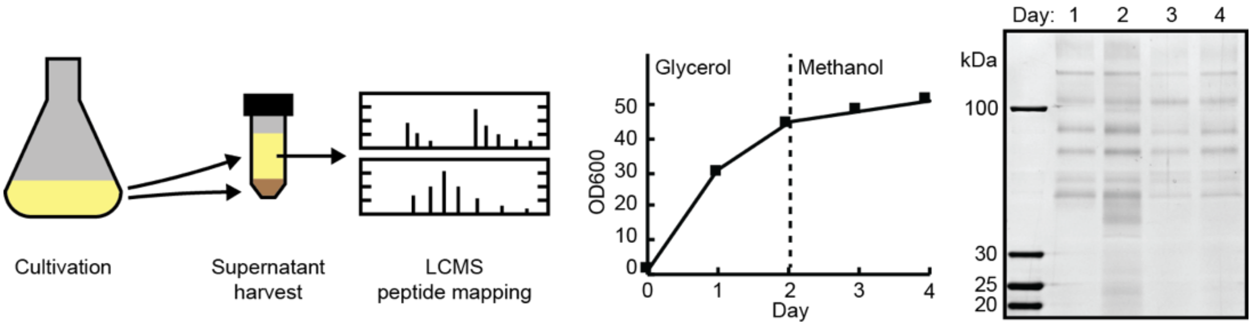


**Fig. S1.** Identification of secreted proteins
Cell growth and harvesting of culture supernatant (left). Representative SDS-PAGE of culture supernatant (right).

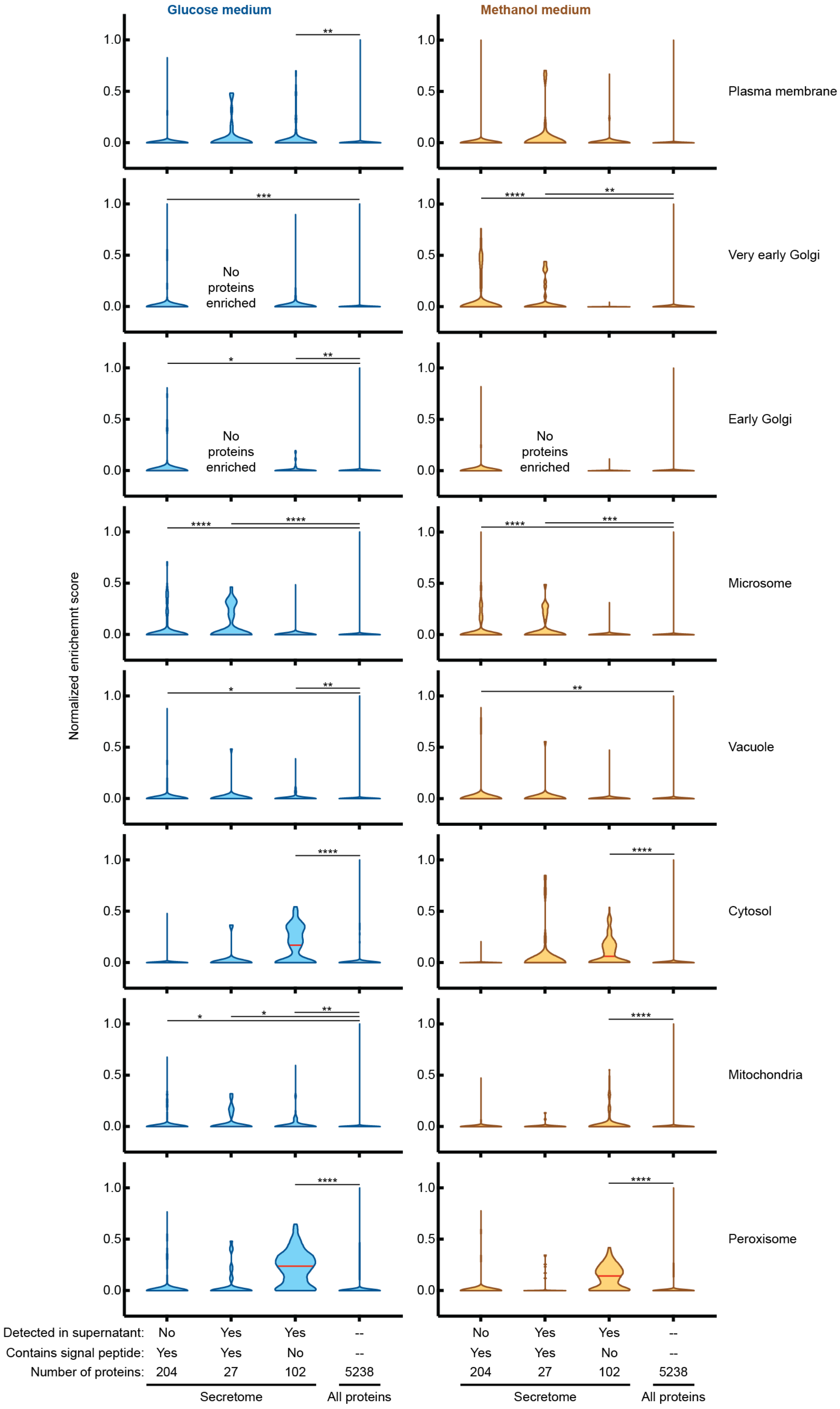
Glucose medium
Methanol medium
Normalized enrichemnt score
1.0
0.5
0.0
Plasma membrane
Very early Golgi
Early Golgi
Microsome
Vacuole
Cytosol
Mitochondria
Peroxisome
No proteins enriched
*
**
***
****
Detected in supernatant: No Yes Yes --
Contains signal peptide: Yes Yes No --
Number of proteins: 204 27 102 5238
Secretome
All proteins

**Fig. S2.** Comparison of the *K. phaffii* secretome to the study by Valli et al. Proteins that were not detected by Valli et al., or that were detected but were below the cutoff, were assigned an enrichment score of 0. Red line represents the median value. Significance was determined by Kruskal-Wallis test with Dunn's multiple hypothesis correction ($p<0.05$*, $p<0.01$**, $p<0.001$***, $p<0.0001$****).

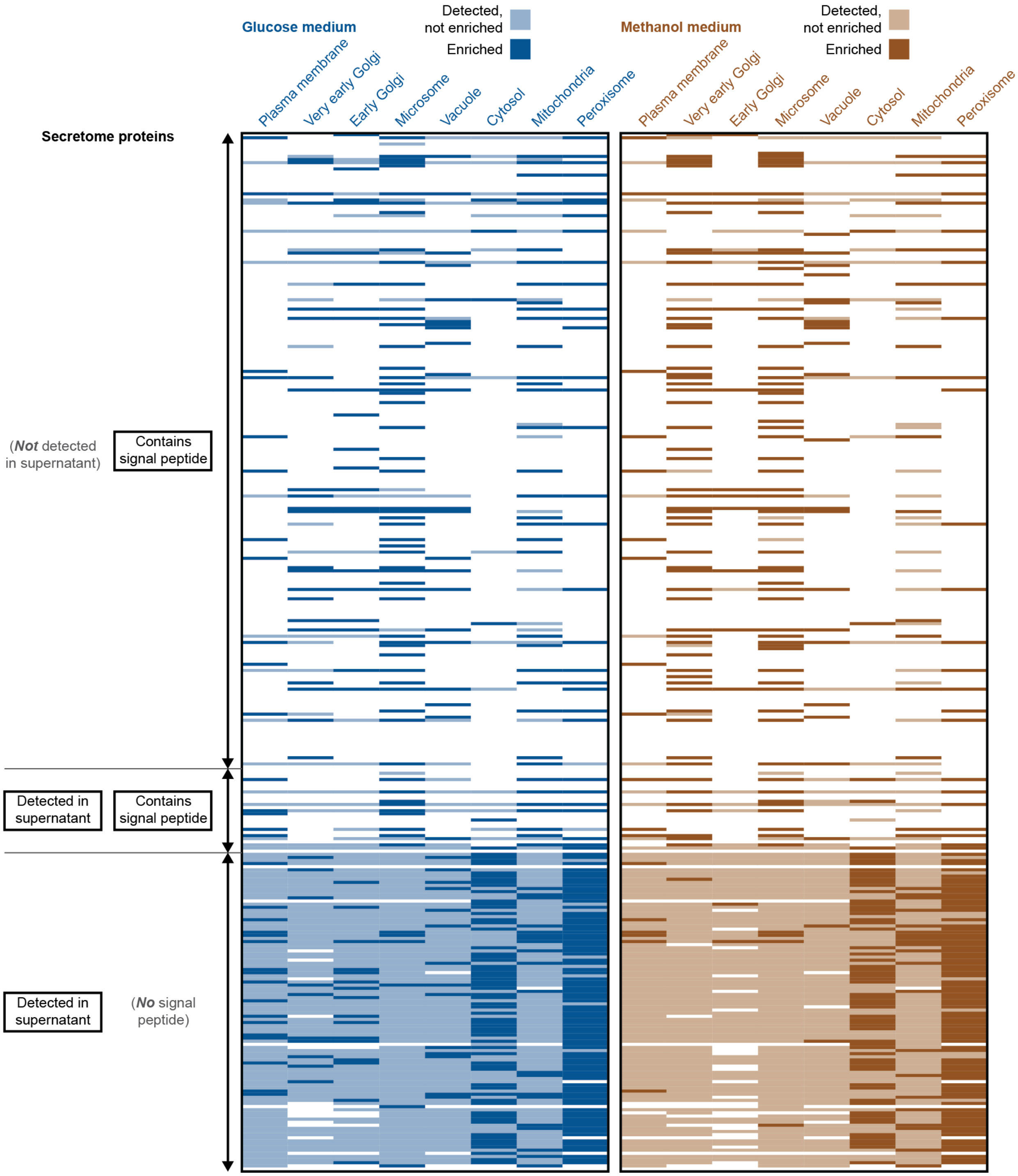


**Fig. S3.** Detection or enrichment of proteins in the *K. phaffii* secretome by Valli et al.

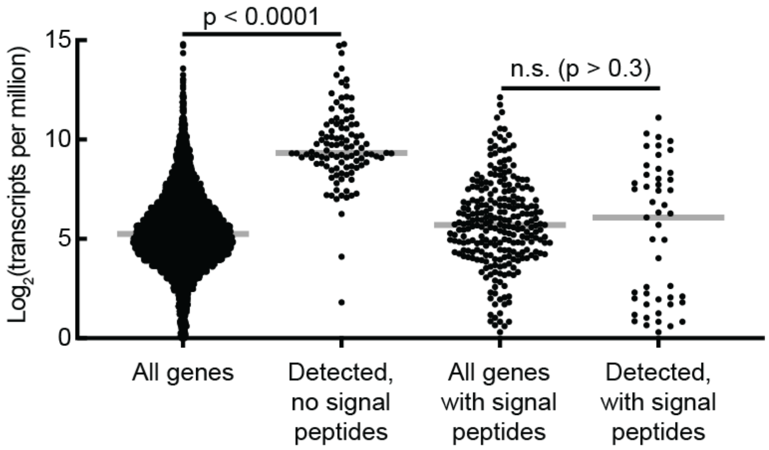


**Fig. S4.** Gene expression from RNAseq of an unedited base strain. Data was adapted from Love, K.R., et al. Genes labeled as "detected" code for proteins that were observed in culture supernatants by LCMS. Signal peptides were predicted by SignalP. Significance was determined by Welch's t-test. Grey bars represent median values.

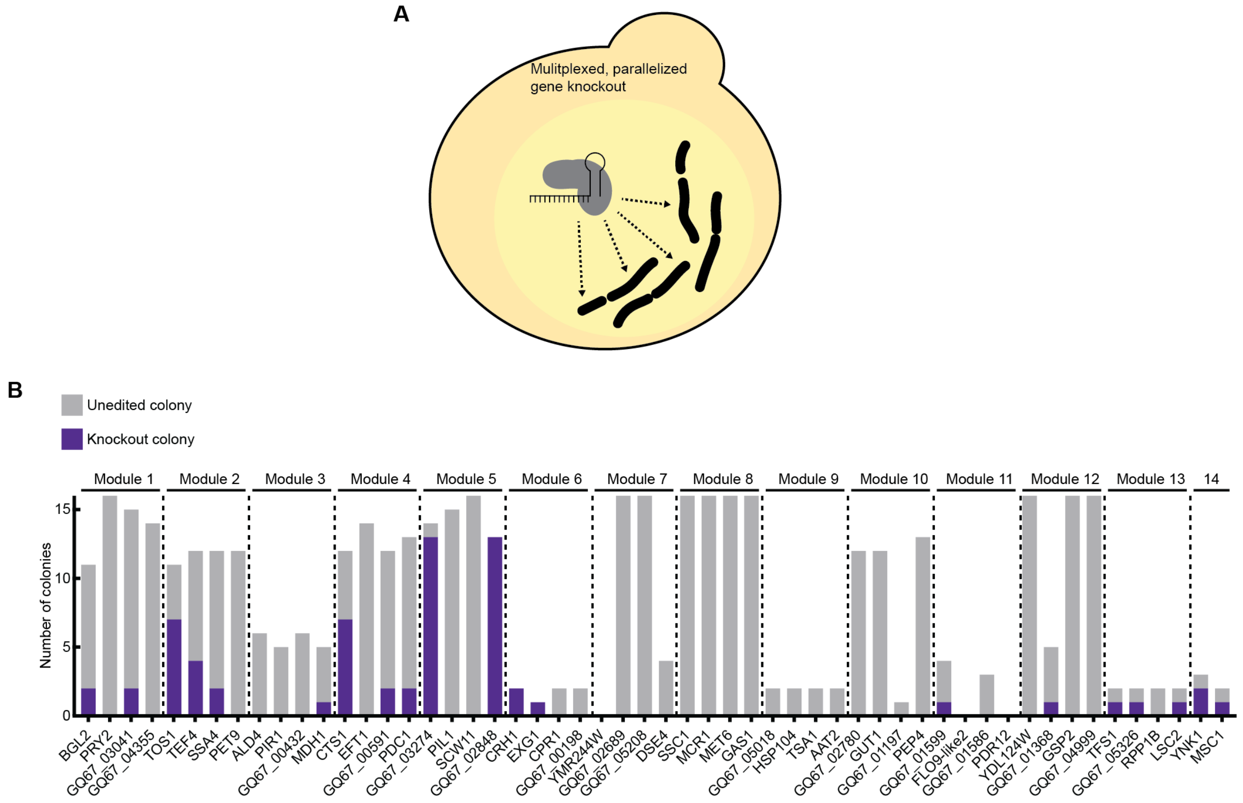


**Fig. S5.** Multiplexed knockout of secretome genes.
A) Schematic of simultaneous knockout of four genes. B) Results of multiplexed knockout screening. Knockout genotypes were determined by Sanger sequencing of each target locus of up to 16 colonies per transformation.
