## Supplement for "CRISPR-Cas9 knockout screen informs efficient reduction of the *Komagataella phaffii* secretome": Supplemental Methods.pdf

**Supplementary methods on creation of a knockout library**

Neil C. Dalvie[1,2], Timothy R. Lorgeree[2], Yuchen Yang[1,2], Sergio A. Rodriguez-Aponte[2,3], Charles A. Whittaker[2], Joshua A. Hinckley[2], John J. Clark[2], Amanda M. Del Rosario[2], Kerry R. Love[1,2]*, J. Christopher Love[1,2]*

[1]Department of Chemical Engineering, Massachusetts Institute of Technology, Cambridge, Massachusetts 02139, United States
[2]The Koch Institute for Integrative Cancer Research, Massachusetts Institute of Technology, Cambridge, Massachusetts 01239, United States
[3]Department of Biological Engineering, Massachusetts Institute of Technology, Cambridge, Massachusetts 02139, United States

### Creation of a *K. phaffii* strain with a genomic copy of Cas9

We used a transient CRISPR-Cas9 plasmid described previously[1] to integrate a permanent cassette for constitutive expression of Cas9 protein into the *K. phaffii* genome at an intergenic region near the gene GQ67_01884 under control of the *K. phaffii* $P_{ENO1}$ promoter (pND386, included in Supplemental Materials).[2]

### Design and construction of knockout libraries

We designed single guide RNAs (sgRNAs) to target all coding sequences in the *K. phaffii* genome. We used CRISPOR[3] to identify up to five sgRNAs within the first 500 bp of each gene with the highest specificity scores.[4] We eliminated sgRNAs that contained 4 bp homopolymers, had a GC content below 20% or above 80%, or were within 5 bp of another higher ranking sgRNA. This resulted in 25425 sgRNAs (Table S3). We then identified the sgRNAs that target the 361 coding sequences in the *K. phaffii* secretome. This resulted in 1777 sgRNAs. We synthesized all 1777 sgRNAs as an oligo pool with cloning handles attached (Genscript). We amplified the library by PCR (primers: CAATTCCCCGTCGCGGAG, CCTTATTTAACTTGCTATTTCTAGCTCTAAAAC) and cloned the library into plasmid pND389 using NEBuilder® HiFi DNA Assembly (New England Biolabs) (backbone primers: GTTTTAGAGCTAGAAATAGCAAGTTAAAATAAGG, CTCCGCGACGGGGAATTG). pND389 is a custom vector for integration of the sgRNA library into the *K. phaffii* genome at an intergenic region near the gene GQ67_02926 (*rpl5*) and is attached in the Supplemental Materials.[1,2] We transformed the HiFi product into 10-beta Electrocompetent *E. coli* (New England Biolabs) according to the manufacturer’s protocol. We propagated the libraries according to a previously described protocol.[5] After extraction of the pooled plasmid library, we linearized the DNA by digestion with PmeI (New England Biolabs) according to the manufacturer's protocol. We transformed 4 µg of linearized DNA into competent *K. phaffii* as described elsewhere,[6] plated cells on a large agar plate (Difco™ YPD Agar, BD), and incubated at 30ºC for two days to allow formation of >$10^5$ colonies. We performed four replicates of transformation into the Cas9(+) strain and into a Cas9(-) (wild type) strain.

**Sequencing pooled knockout libraries**

We scraped all colonies from each individual transformation plate into a pooled cell bank and extracted genomic DNA. We used nested PCR to amplify the genomic sgRNA cassette and add sequencing handles and barcodes (Inside primers: AATCTTGAAGAAGCCACCCGTCGCGGAGC, GTGACTGGAGTTCAGACGTGTGCTCTTCCGATCTACCTTTAGTACGGGTAATTAACGACAC; Outside primers: AATGATACGGCGACCACCGAGATCTACACAATCTTGAAGAAGCCACCCG, CAAGCAGAAGACGGCATACGAGATNNNNNNGTGACTGGAGTTCAGACGTG). We sequenced the amplicon library on an Illumina HighSeq2000 (40 bp, single end) with a TruSeq i7 index primer and a custom primer for the genomic sgRNA sequence (AATCTTGAAGAAGCCACCCGTCGCGGAGC). We also sequenced the original plasmid library. We tabulated the number of read counts of all sgRNAs in each sample. We observed fewer sgRNAs in the Cas9(+) library than in the Cas9(-) library, which suggested that sgRNAs that target essential genes may be depleted from the library when Cas9 is expressed.

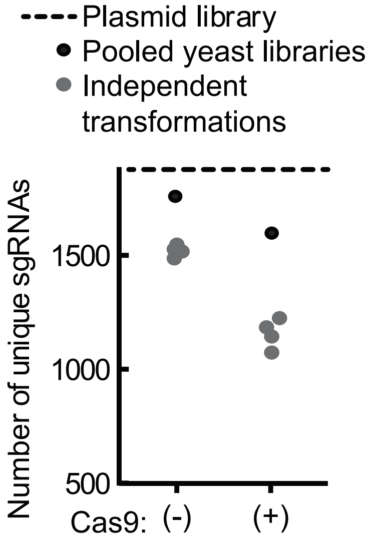


*Fig. SM1. Number of unique sgRNAs detected in amplicon sequencing with at least 100 read counts from independent transformations of the CRISPR-Cas9 knockout library. Black points represent the union of all independent transformations in each strain background.*

**Assessment of sgRNA copy number**

To approximate the multiplicity of transformation, we extracted gDNA from 16 transformants (from individual colonies) and included these samples in amplicon sequencing. We observed a single dominant sgRNA for 14 of the clones, and two dominant sgRNAs for 1 of the clones. We concluded then, that most members of the yeast knockout library had received a single copy of the sgRNA integration cassette, and therefore a single sgRNA.

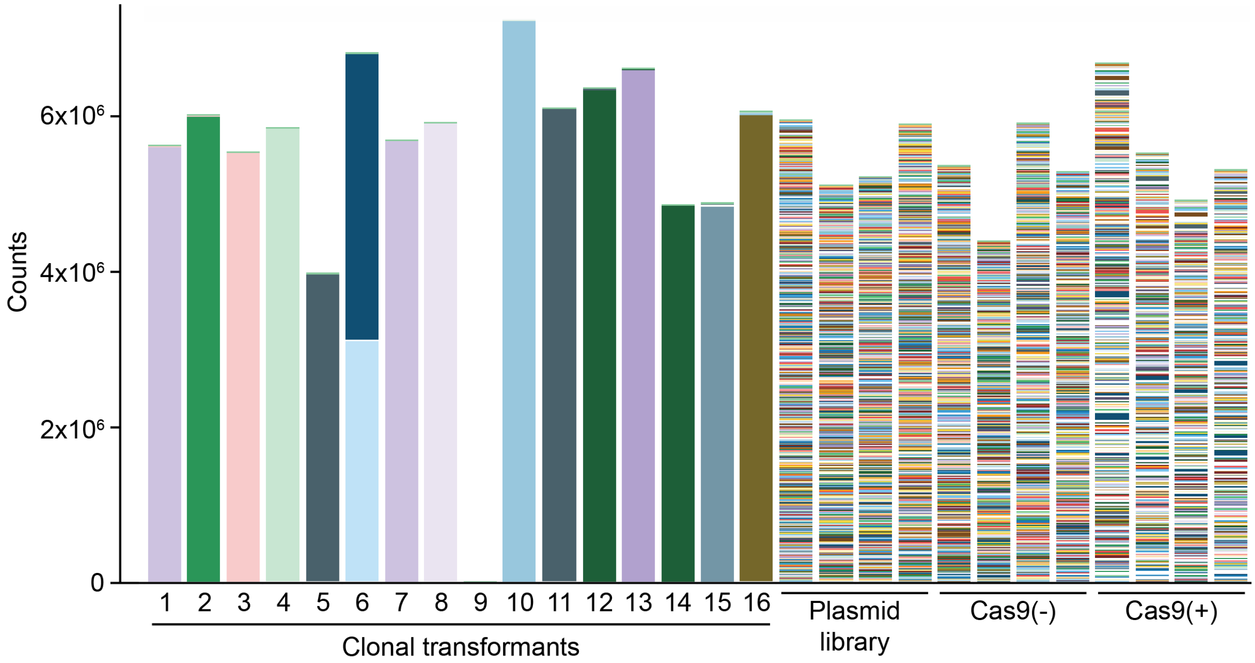


*Figure SM2. Bar plot of unique sgRNAs in each sequenced colony or pooled library. Each color represents one guide RNA.*

**Analysis of pooled knockout libraries**

We calculated the relative abundance of each sgRNA within each sample. The relative abundance (A) of sgRNA (i) with counts (C) in replicate (j):

$$A_{i,j} = \frac{C_{i,j}}{\sum_i C_{i,j}}$$

We then calculated the log fold change in abundance of each sgRNA between the pooled yeast library and the original plasmid library and averaged over the four replicates.

$$lfc_{i,j} = \log_2 \frac{A_{i,j,yeast}}{A_{i,plasmid}}$$

$$lfc_i = \frac{lfc_{i,j}}{\sum_j lfc_{i,j}}$$

We then evaluated the difference in log fold change of each sgRNA between the Cas9(+) and Cas9(-) yeast libraries.

$$LFC_i = lfc_{i,Cas9(+)} - lfc_{i,Cas9(-)}$$

We observed that more sgRNAs were differentially represented in the Cas9(-) strain, which may indicate that these genes are depleted in the Cas9(+) strain. All abundance and fold-change calculations are reported in Table S3.

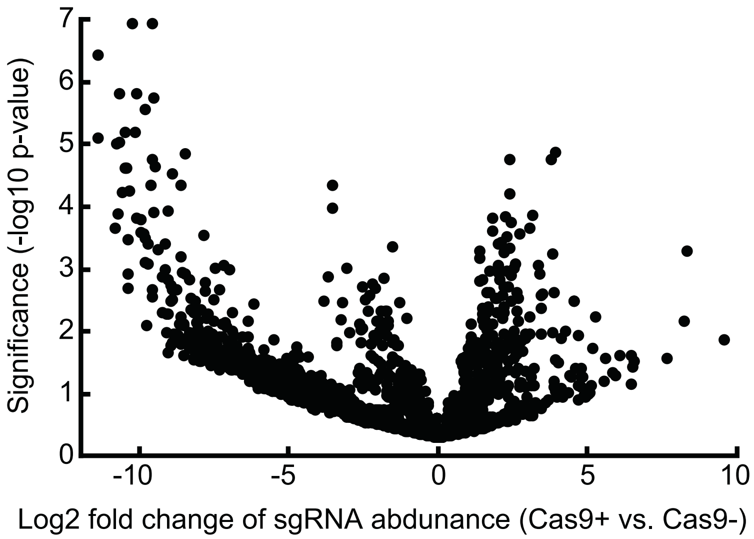


*Fig. SM3. Difference in $\log_2$ fold change of sgRNA abundance between the Cas9(+) strain and the Cas9(-) strain. Each point represents one sgRNA. Significance was determined by a 1-tailed heteroscedastic t-test across 4 experimental replicates.*

Finally, we performed a one-sided Kolmogorov-Smirnov (KS) test on the LFC of all sgRNAs for each gene. We defined essentiality of each gene as the negative $\log_{10}$ of the KS test p-value. R-code for the KS-test is included in the Supplemental Materials.
